# Endless Conflicts: Detecting Molecular Arms Races in Mammalian Genomes

**DOI:** 10.1101/685321

**Authors:** Jacob C. Cooper, Christopher J. Leonard, Brent S. Pedersen, Clayton M. Carey, Aaron R. Quinlan, Nels C. Elde, Nitin Phadnis

## Abstract

Recurrent positive selection at the codon level is often a sign that a gene is engaged in a molecular arms race – a conflict between the genome of its host and the genome of another species over mutually exclusive access to a resource that has a direct effect on the fitness of both individuals. Detecting molecular arms races has led to a better understanding of how evolution changes the molecular interfaces of proteins when organisms compete over time, especially in the realm of host-pathogen interactions. Here, we present a method for detection of gene-level recurrent positive selection across entire genomes for a given phylogenetic group. We deploy this method on five mammalian clades – primates, mice, deer mice, dogs, and bats – to both detect novel instances of recurrent positive selection and to compare the prevalence of recurrent positive selection between clades. We analyze the frequency at which individual genes are targets of recurrent positive selection in multiple clades. We find that coincidence of selection occurs far more frequently than expected by chance, indicating that all clades experience shared selective pressures. Additionally, we highlight Polymeric Immunoglobulin Receptor (PIGR) as a gene which shares specific amino acids under recurrent positive selection in multiple clades, indicating that it has been locked in a molecular arms race for ∼100My. These data provide an in-depth comparison of recurrent positive selection across the mammalian phylogeny, and highlights of the power of comparative evolutionary approaches to generate specific hypotheses about the molecular interactions of rapidly evolving genes.

## Introduction

Across all life forms, a repertoire of highly conserved core genes allows cells to function with extreme consistency. Most of the time, mutations in these genes are deleterious to the fitness of an individual, and therefore do not persist. However, rarely mutations cause changes to a gene which increase the fitness of the individual. If these mutations continue to convey fitness advantage to individuals, they will eventually rise in frequency in a population – this is referred to as positive selection. Perhaps because they are at the interface between a host and a pathogen, or because they have a direct impact on fertility, some genes are a repeated target of positive selection – this is referred to as recurrent positive selection (RPS). Despite the importance of RPS in identifying the fastest and most dynamic evolutionary processes, it remains unclear how common selective pressures might drive RPS in specific genes across diverse groups of animals.

Detection of RPS has proven to be a particularly powerful tool for dissecting molecular interfaces where hosts and pathogens interact [1–7]. The recurrent evolution at these host-pathogen interfaces are often referred to as a molecular arms races, where the profound fitness consequences for both the host and pathogen result in RPS [8]. When comparing multiple species, arms races that occur in the protein-coding region of genes leave signals that can be detected by measuring the rate of amino acid fixation against the rate of neutral change. Analyzing the rapidly evolving regions of protein-coding sequences has illuminated portions of genes that are important for hosts and pathogens to interact, on what would otherwise be complex binding interfaces. When coupled with functional studies, these approaches have also furthered our understanding of pathogen tropisms, where rapid diversification of host genes that evolve under RPS often results in highly specific species interactions. Therefore, determining the genome-wide portfolio of RPS genes is important for understanding the evolutionary history of a clade of species and as a method for identifying and understanding the mechanisms of host-pathogen interactions.

There have been multiple efforts to scan genomes for RPS in primates and other mammals, both for understanding selective pressures acting on immune genes and for the discovery of new rapidly evolving processes, [9– 13]. The main pattern revealed by these studies is that immune genes are much more likely to be rapidly evolving than the rest of the genome, leading to further analyses that have suggested pathogens to be the main drivers of rapid evolution [14]. These studies have been limited, however, by the use of only a fraction of currently available mammalian genomes, thereby decreasing the power to detect RPS [15]. Therefore, it is likely that many cases of RPS have gone undetected thus far. Furthermore, these studies focus on understanding evolution in single clades of mammals, which all experience distinct sources of selective pressure. Therefore, it remains unclear whether the patterns of RPS observed in primates are broadly shared among other mammals, or whether the repertoire of rapidly evolving genes is largely unique in each clade.

Here, we design a computational pipeline to execute the most widely-used program for detecting recurrent positive selection at a genome-wide scale. We deploy this pipeline to scan for recurrent positive selection in five different clades of mammals, including primates, two clades of rodents, caniforms, and bats. We find many novel examples of recurrent positive selection across all these clades, and provide the results as a resource for others examining specific genes of interest. We ask how frequently single genes show signs of recurrent positive selection across multiple clades and find that there is a much higher coincidence of recurrent positive selection than to be expected by chance. Finally, we examine specific molecular interfaces that have been under recurrent positive selection in multiple clades. As an example, we highlight Polymeric Immunoglobulin Receptor (PIGR) as a gene that has seen recurrent positive selection in the same amino acids across multiple clades in mammals, suggesting that it is an import component of a yet undescribed molecular arms race. Our results will inform future studies focused on recurrent positive selection, and provide many new examples that will help to understand the evolutionary forces that shape the genome.

## Results

### Recurrent positive selection in five clades of mammals

To compare recurrent positive section across mammals, we identified five clades that have a sufficient number of genomes to conduct this analyses: Primates (humans, apes, monkeys), Murinae (mice, rats), Cricetidae (deer mice, hamsters), Chiroptera (bats), and Caniformia (dogs, bears, seals). We also included a sub-sampling of the primates clade that was more comparable to the other clades in the number of genomes used. We gathered all publicly available genomes for each species in each clade (Supplemental Table 1) and used gene sequences from the best-annotated genome in each clade as the reference [16]. We then identified homologous gene sequences for each gene that has a 1:1 orthologous relationship within its clade, as this analysis is easily confounded by paralogous sequences. We aligned these sequences and tested them for evidence of recurrent positive selection using Phylogenetic Analysis by Maximum Likelihood (PAML) [17]. For this test, we first measured the log-likelihood difference between Model 7 (no positive selection) and Model 8 (positive selection). A significant difference between the fit of these two models to the distribution of the rate of codon evolution in a gene is evidence that the gene has undergone recurrent positive selection. If the differences between these two models were significant, then we compared Model 8 to Model 8a (neutral evolution), to test against a scenario of neutral evolution that can be missed by Model 7. In our analysis, we did not require that every gene be identified in all species to complete the analysis. Instead, we required that 90% of the gene be identified in at least four species. We report the distribution of the number of species used for each clade (Supplemental Figure 1).

We find that recurrent positive selection is pervasive in all clades, as has been reported for primates and mice in previous work [11,12]. In all clades that we analyzed, between 1.3% (Murinae) and 6.5% (Chiroptera) of the genes analyzed showed a signature of recurrent positive selection by the M8-M8a test after multiple testing correction (Table 1). Phylogenetic distance can be a confounding factor for PAML, as long branch lengths make estimations of dN and dS inaccurate. To retroactively confirm that all the clades that we used had an appropriate phylogenetic distance for PAML, we calculated the maximum dS for each gene run in each clade (Supplemental Figure 1). We find that the average maximum dS is close to 0.25 for all clades, confirming that all of these clades are appropriate for PAML.

**Table 1.**
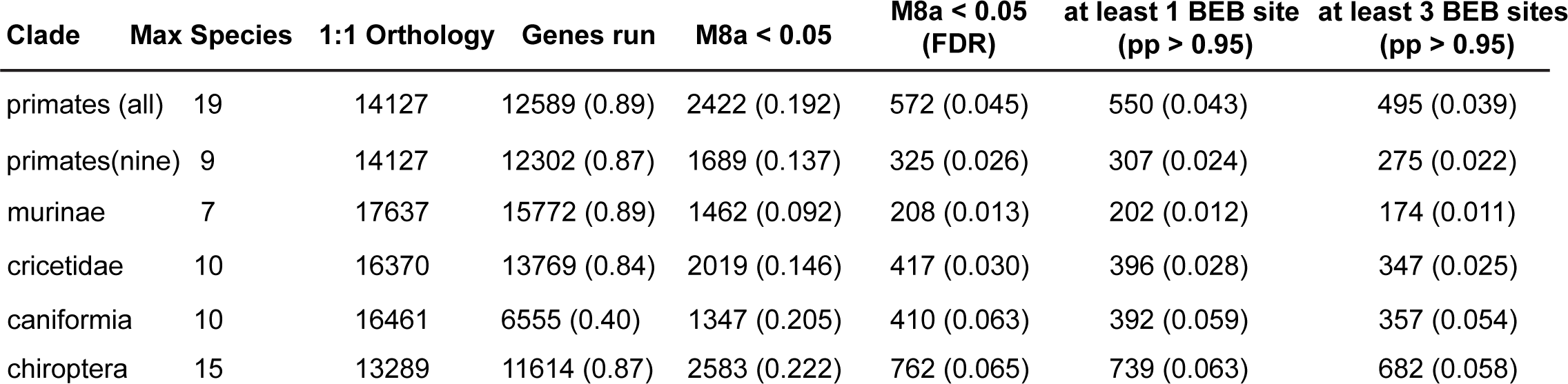
Recurrent Positive Selection in Five Mammalian Clades.

Another form of evidence for recurrent positive selection is the toggling at specific amino acid positions in multiple species in a phylogeny. These sites are of particular interest because they are the most likely to represent the interface of a molecular arms race, which has been demonstrated experimentally numerous times. To identify these sites, we calculated the Bayes Empirical Bayes (BEB) posterior probability (pp) that the state changes in each site throughout the gene were driving the signature of recurrent positive selection and considered a site to be significant at a BEB pp > 0.95. Again, we find that all clades have several hundred genes with at least one or more of these sites (Table 1). In our subsequent analyses, we use genes that had significant results for the M7-M8 and M8-M8a tests, and have one amino acid under selection with a BEB pp > 0.95 as our list of positively selected genes.

### Patterns of Pairwise Recurrent Positive Selection in Mammalian Clades

We next investigated the coincidence of recurrent positive selection between clades. First, we made pairwise comparisons of positively selected genes across all clades (Figure 2A) and then compared the frequency of overlapping genes against the frequency expected by chance. A greater overlap of rapidly evolving genes than expected by chance indicates that the evolutionary forces that drive rapid positive selection are shared even across multiple clades. We find that all clades have a greater number of coinciding genes under recurrent positive selection than expected by random chance, with the greatest coincident score between Murinae and Cricetidae (Figure 2B). These two clades are closely related and have a high degree of life-history similarity, indicating that these two clades might share similar selective pressures that drive recurrent positive selection. We do not find any comparison with fewer coinciding genes than expected, indicating that the evolutionary pressures that drive diversifying selection between clades are not entirely distinct.

**Figure 1.**
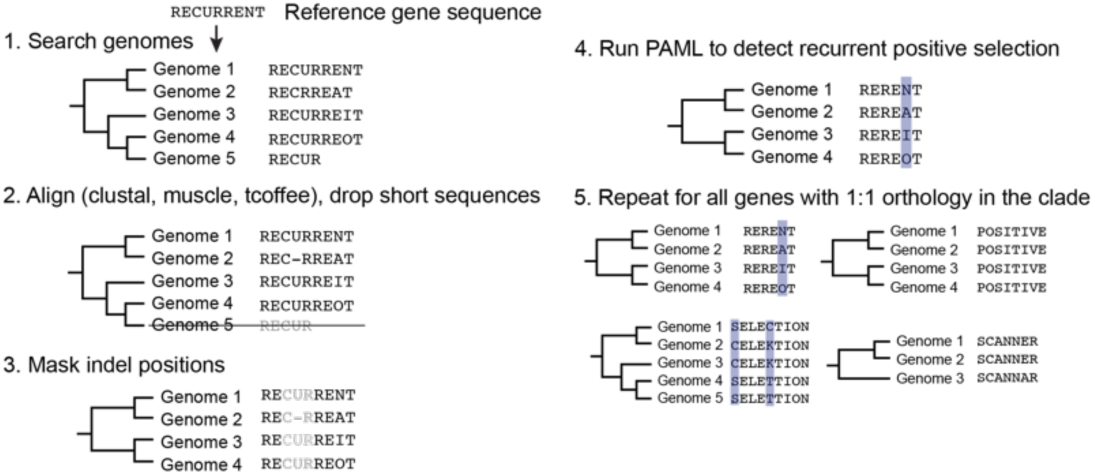
Scanning for recurrent positive selection. Our pipeline is designed to run PAML on the scale of an individual gene, and can be easily scaled to the genomic scale. Here we use a toy example of the word “recurrent” to illustrate the process. 1) Search all species in a clade for gene sequence using the reference gene amino acid sequence. 2) Align the genes for each species using ClustalO, muscle, and Tcoffee. Drop any sequences that are less than 90% of the reference gene length (less is kept here for clarity). 3) Using the consensus alignment, mask indel codons and +/-1 of indel codons. 4) Use PAML to detect recurrent positive selection. 5) Scale this process to analyze every gene in the clade that has a 1:1 orthology relationship throughout the clade.

**Figure 2.**
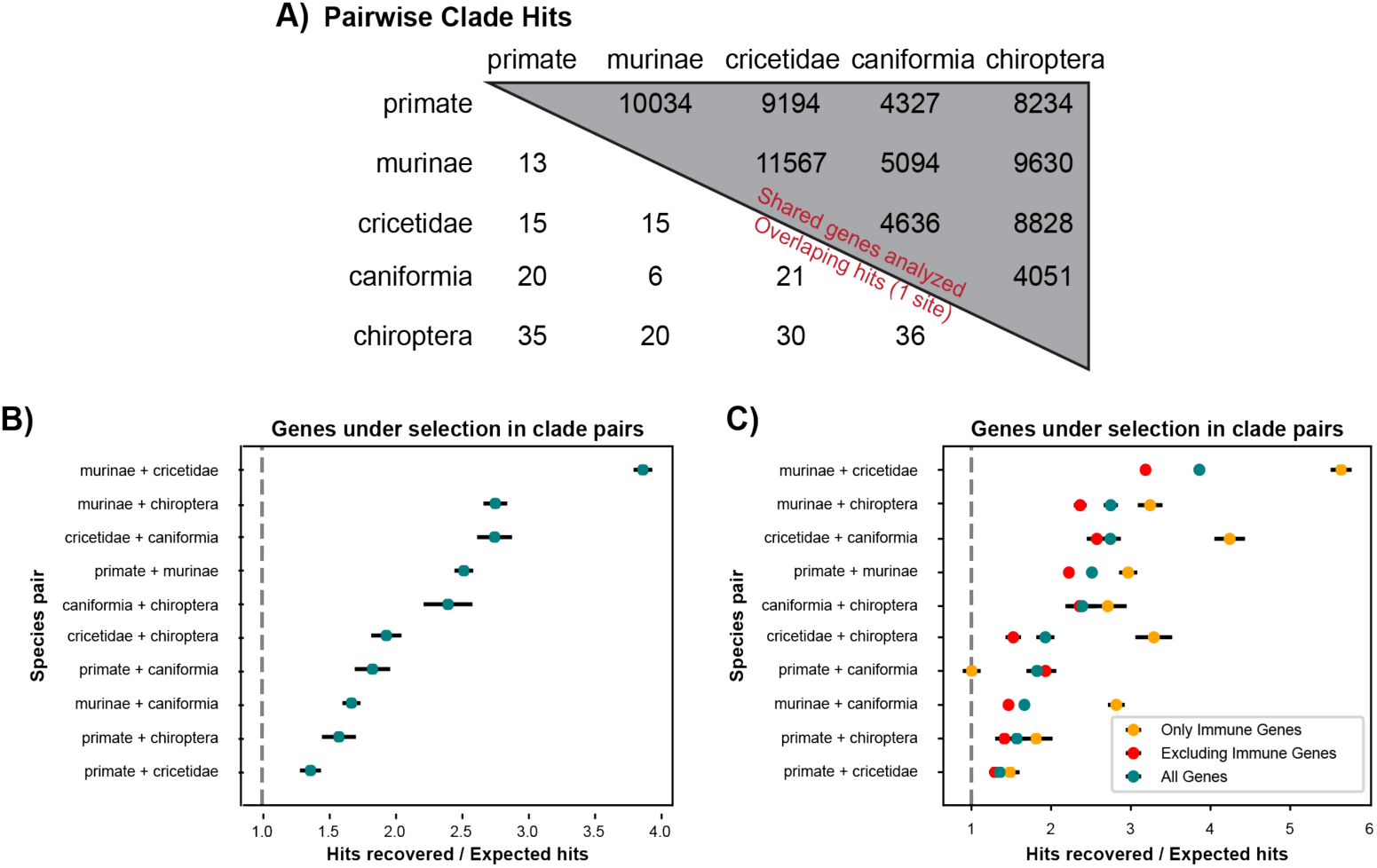
Pairwise clade analysis. Genes that scanned positive for recurrent positive selection in two clades occur more frequently than expected. A) Table with the number of genes analyzed for each comparison (top half) and the number of hits between each comparison (bottom half). B) Rate of pairwise hits for recurrent positive selection vs. the rate of hits expected by chance. The 95% confidence interval is given as a black bar around each point. Every comparison had a greater ratio of hits recovered than expected, with the neutral value represented by the dashed line. C) Contribution to the signal in B from immune genes (red) and non-immune genes (yellow). The data from B is re-plotted for comparison (blue).

All organisms are constantly under attack from pathogens in the environment, and previous analyses of rapidly evolving genes have indicated immune function as a common driver of diversifying selection [14]. To examine the influence of immune function in our test for coincident recurrent positive selection, we segregated the list of rapidly evolving genes into those that have roles in infection - and those that do not - based on KEGG pathway annotation (Figure 2C). We then re-analyzed the coincidence of recurrent positive selection in these two subsets of the data. We detected recurrent positive selection with a higher degree of coincidence than by chance alone for both immune genes and non-immune genes. In this analysis, there is a general trend that the set of immune genes has a greater magnitude of deviation from the expectation of hits recovered than the set of non-immune genes (6/10 comparisons). We find only one case where the opposite trend is true, and several cases where no distinction can be made (3/10). This analysis demonstrates that specific immune genes are targets of recurrent positive selection repeatedly over multiple clades. However, our data clearly show that immune genes do make up the entirety of the signal for recurrent positive selection between clades.

### Recurrent current positive selection of specific amino acid positions in multiple clades

Genes that experience recurrent positive selection in more than two lineages may be interesting candidates to study because they represent truly long-term (or constantly restarting) arms races in the mammalian lineage. Beyond comparison of two clades, however, we do not have sufficient power to make clear conclusions about the rate of the expected and observed coincidence of recurrent positive selection. Still, there are 20 genes with shared signatures of recurrent positive selection in three clades in our data, and two genes (*PIGR* and *CD72*) with signatures of recurrent positive selection in four clades. We did not identify any genes with recurrent positive selection in five clades (Table 2). Of the multi-clade genes we did identify, most of them represent novel cases of detecting any form of recurrent positive selection; previous characterization of positive selection in any lineage only exists for *SERPINC1, TSPAN8, SAMD9*L [12,18–20].

**Table 2.**
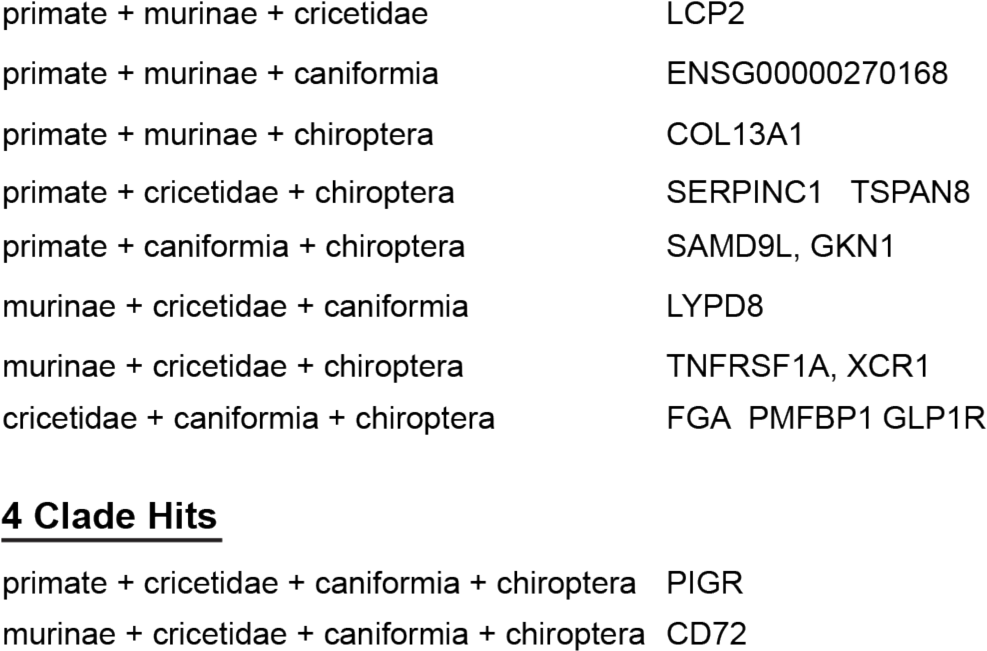
Genes with signatures of recurrent positive selection in three and four clades

|  |  |
| --- | --- |
| primate + murinae + cricetidae | LCP2 |
| primate + murinae + caniformia | ENSG00000270168 |
| primate + murinae + chiroptera | COL13A1 |
| primate + cricetidae + chiroptera | SERPINC1 TSPAN8 |
| primate + caniformia + chiroptera | SAMD9L, GKN1 |
| murinae + cricetidae + caniformia | LYPD8 |
| murinae + cricetidae + chiroptera | TNFRSF1A, XCR1 |
| cricetidae + caniformia + chiroptera | FGA PMFBP1 GLP1R |
| 4 Clade Hits |  |
| primate + cricetidae + caniformia + chiroptera | *PIGR* |
| murinae + cricetidae + caniformia + chiroptera | CD72 |

To ask if the same interfaces of the recurrently selected genes have been exposed to selection in multiple clades, we analyzed the positioning of amino acids under recurrent selection in each of the 15 genes that show recurrent selection in 3 or more clades (Supplemental Figure 3). From this analysis, it is clear that several genes have clusters of amino acids that are under recurrent positive selection in multiple clades, implying that a single molecular interface is the recurrent target of some selective pressure. This pattern is most obvious in *ENSG00000270168, TSPAN8*, and *PIGR*. The greatest density of selected sites in multiple clades appears to fall in *PIGR*.

PIGR (Polymeric Immunoglobulin Receptor) is a receptor on the basal surface of mucosal epithelial cells that binds to dimeric IgA and pentameric IgM and transports them to the apical surface [21]. In our data, we find evidence that is under recurrent positive selection in Primates, Cricetidae, Caniformia, and Chiroptera. In Murinae, it appears that the reference sequence for *PIGR* is truncated by ∼560 amino acids, and we did not analyze the entire sequence. Therefore, it is unclear if recurrent positive selection in this gene extends to Murinae as well.

To better understand the spatial distribution of the codons under selection in this gene, we plotted the changes on a recently solved structure of PIGR [22] (Figure 3). PIGR contains five immuno-globulin domains (D1-D5) [23]. We find that the sites under selection in *PIGR* are heavily enriched on the outfacing section of domain D2. Though *PIGR* has been studied for over 20 years [21], little is known about the specific function of the D2 domain. Evidence suggests that it plays a role in the transport of IgM but not IgA [24], and as of yet, there has been no evidence of a role for this domain in the innate pathogen binding actions of *PIGR* [25]. Our data suggest that the outer face of the D2 domain participates in a host-pathogen type evolutionary arms race across most mammals, indicating that it has an important role in immune function that is yet to be discovered.

**Figure 3.**
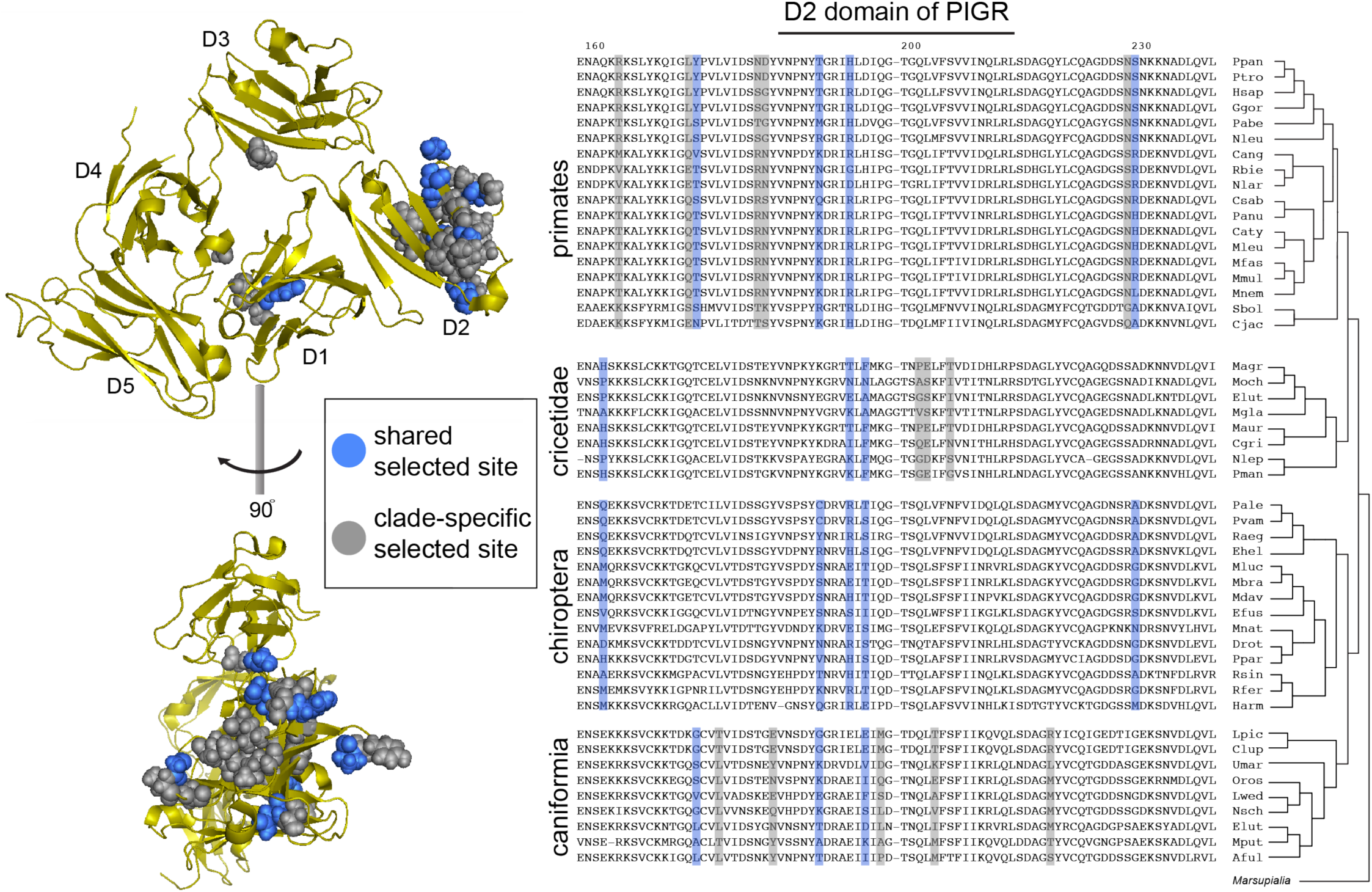
PIGR evolves under recurrent positive selection in multiple clades of mammals. The crystal structure of PIGR is plotted in yellow on the left, with the immunoglobulin domains annotated as D1-D5. Sites under selection are shown as globular residues as either recurrently selected in multiple clades (red) or recurrently selected in just one clade (grey). The structure is presented in two orientations, so that the domains are visible (top), and rotated 90 degrees clockwise so that the arrangements of the sites are visible (bottom). The amino acid alignment on the right shows the D2 region of PIGR for each species analyzed. Selected sites are denoted using the same color scheme as the structure on the left. Species names are given as a four-letter code (ex: *Homo sapiens* – *Hsap*), with a cladogram of their relationships on the far right.

## Discussion

The modern age of genomics has brought with it many whole sequenced genomes over a broad taxonomic diversity, allowing evolution to be studied in a way that was not possible before. In this study, we sought to leverage this deep and diverse resource to analyze patterns of recurrent positive selection in five different clades of mammals. We designed a pipeline for increasing the throughput of the Phylogenetic Analysis by Maximum Likelihood (PAML) [17] so that we could deploy this analysis on a genome-wide scale multiple times over. Subsequently, we asked if our analysis detected instances of recurrent positive selection in the same genes over multiple clades. We found instances of recurrent positive selection in the same genes far more often than expected by chance in every pairwise comparison, indicating that there is an excess of genes that are frequently the targets of positive selection. We narrowed our comparison to some of the most frequently recurrently selected genes and identified a single molecular face of *Polymeric Immunoglobulin Receptor* (*PIGR*) that has been the target of recurrent positive selection in mammals for nearly 100 million years.

In developing our method, we identified several sources of error that would lead to false positives and took care to eliminate them. First, PAML is highly sensitive to insertions and deletions (indels), as even slight misalignment of sequences can easily replicate the signature of recurrent positive selection [26,27]. To circumvent this problem, we developed a strategy of aligning each gene with three different aligners, then only retaining sites where all aligners agreed on the alignment. This is similar to previous approaches that have used a post-hoc test to compare the results from multiple runs of PAML using different aligners [28], but our method avoids creating false negatives when one section of a gene contains indels, but another section of the gene has a robust signature of positive selection.

This study follows on a history of other studies which have analyzed recurrent positive selection in primates, each time improving the analysis as new resources become available. In our study, our rate of detection for positively selected genes in primates falls in line with previously observed rates - 4.5% in our analysis vs. 1%-10% in previous work [9–12,26,29,30]. In our data, every clade that we analyzed had rates of positive selection in this range. Like previous studies, we consider our detection method to be extremely conservative, and much more likely to identify false negatives than false positives. It is therefore likely that this is an underestimate of recurrent positive selection in most of these cases. Our comparison between clades then relies on the fact that we used the same method of detection in every case, not that our data represents a ground truth about recurrent positive selection.

Encouragingly, our data set identifies several previously studied examples of recurrent positive selection in at least one clade: for example, *OAS1* [31], *PKR* [4], *TFRC* [3], *IZUMO1* and *IZUMO4* [32], *PARP4* and *PARP15* [6], *CATSPER 1-4, D, G*, and *E* [33], *RNASEL* [34], *ZP3* [35], and *CGAS* [5]. We compared our results with a list of genes representing the most heavily enriched genes for selected sites from a previous genome-wide scan in primates and found that *MUC13, NAPSA, PTPRC, APOL6, MS4A12, SCGB1D2, PIP, CFH, RARRES3, OAS1*, and *TSPAN8* were detected under recurrent positive selection in at least one clade, while our analysis missed *PASD1, CD59*, and *TRIM* (11 / 14 genes) [12]. Given the differences in the methodology of the two studies, we believe that this represents a high degree of replication of signals for recurrent positive selection, which has been difficult to obtain in previous work.

Our analysis is the first to scan for recurrent positive selection over the entire genomes of multiple mammalian clades. We used this new set of data to ask how often we detected recurrent positive selection in the same gene in multiple clades. In our pairwise comparison between clades, we detected far more coincident recurrent positive selection than expected by chance. Recent work has implicated immune genes as the main targets of positive selection in multiple lineages [14,30], raising the possibility that immune genes drive the signal we detect. When we asked how immune genes contributed to this trend, we found that immune genes have a higher rate of coincidence of recurrent positive selection than the set of all other genes, though they did not explain the entire signal. Combined with previous work, our data suggest that while immune genes as a class are more frequently the target of recurrent positive selection, specific immune genes are not the only genes to experience recurrent positive selection in multiple clades more often than expected. It is difficult to say whether the few examples of recurrent positive selection that we find in three clades and four clades constitute a greater rate than expected because the expectation of the number of genes to recover from these comparisons is too low to be biologically meaningful.

However, our data is appropriate for analyzing specific molecular interfaces that might be under recurrent positive selection in multiple clades. Recurrent evolution on three-dimensional protein interfaces are signs of molecular arms races, thought to be waged over interactions at that interface. Identifying these interfaces has been a fruitful approach in understanding the otherwise very complex interactions between host proteins and pathogen-derived factors that might try to interfere with their functions, often identifying specific amino acids or pockets of binding that are crucial for the interaction. In our data, we identify *PIGR* as having a specific molecular face that has been the target of recurrent positive selection in multiple clades. *PIGR* contains five extracellular immunoglobulin domains, D1-D5 [23], and most of the sites under recurrent selection in both one clade and multiple clades fall in the D2 domain.

*PIGR* was first characterized more than two decades ago and has been the subject of thorough study. *PIGR* is primarily responsible for transporting dimeric IgA and pentameric IgM from the basal to the apical surface of mucosal epithelial cells [21]. *PIGR* is cleaved and secreted from the apical surface while binding to IgA and IgM, where it is protects secreted IgA and IgM from proteolytic degradation [25]. *PIGR* has some native anti-bacterial activity [25,36], but this function does not rely on D2. Alternatively, *PIGR* is endocytosed from the cell surface, and several pathogens exploit this feature to gain entry into mucosal cells [37,38]. *Streptococcus pneumoniae* accomplishes this via the choline-binding protein SpsA, which does bind to fragments of the D2 domain [38]. It is possible that the rapid evolution of the D2 domain is primarily to avoid this or other pro-pathogen interactions of *PIGR*. Given the prevalence of this signature of RPS, further investigation of *PIGR*’s D2 domain may provide general insight into a very generalizable route of attack by pathogens.

As opposed to our finding that recurrent positive selection is more common in specific genes than expected, it is worth mentioning that by quantity, most of the recurrent positive selection that we detect is still clade-specific. Many of these hits represent novel examples of recurrent positive selection, which may prove interesting to study for reasons specific to the biology of those clades. Understanding the patterns of recurrent positive selection in the genome has been a useful pursuit both for understanding patterns of evolution and for studying health relevant host-pathogen molecular arms races. Our results add to this growing body of work by contributing data from multiple clades of mammals and point to examples where rapid evolution is the norm rather than the exception. There is a wide breadth of evolutionary history to study, and our results will provide context for future studies looking to analyze ecologically or medically significant instances of recurrent positive selection.

## Methods

### Data acquisition and distribution

Genome sequences for all species were obtained from the National Center for Biotechnology Information (NCBI). Acquisition numbers, taxonomic IDs, and the web link to each genome are presented in Supplemental Table 1 [9,39–67]. The coding DNA sequences for all genes in each reference genome was obtained from Ensemble (https://ensembl.org/index.html). The full phylogenetic relationships for each clade (Supplemental Figure 1) was constructed from the NCBI Taxonomy Browser. To facilitate use for future projects, we have assembled all code and dependencies into a python package (https://github.com/jcooper036/corsair).

### Sequence Curation

To identify gene sequences throughout each clade, we developed a search method that utilizes the CDS sequence of one well-annotated species in that clade. This method was slightly modified to be a whole genome scale version of a previous method we used to study positive selection for a much smaller group of genes [33]. We started with a list of all protein-coding CDS sequences for our reference species. For each sequence, we searched for the homolog of each sequence by first using tBLASTn [68] to identify the genomic scaffold where the sequence resided in each species followed by exonerating [69] to generate a CDS model. We then removed any sequences that did not represent at least 95% of the total length of the gene. As stop codons can represent pseudogenization, we removed sequences that contained stop codons more than 5% of the distance from the end of the gene (if there was a stop codon in this range, we clipped away everything that came after it). We aligned the protein translations of these sequences with Clustal Omega [70], T-coffee [71], and Muscle [72]. Because PAML can be easily confounded misalignment caused by insertions and deletions, we compared all three alignments and only kept positions that were agreed upon by all three aligners, while additionally trimming one codon on either side of an insertion or deletion. We then determined the phylogenic relationship for the remaining species using the Phylo module of Biopython [73].

### Identification of positively selected sites

To test for recurrent positive selection in a given gene, we used a Phylogenetic Analysis by Maximum Likelihood (PAML) [17], specifically testing between two different models of codon evolution – M7, which fits a beta distribution to the frequency of dN/dS by site but limits the max of the distribution to dN/dS = 1, and M8, which is a similar model except that there is no max on the distribution. A better fit to M8 than M7, as determined by a log-likelihood ratio test with a p-value < 0.05, indicates that the evolution of that gene for those sequences is best explained by a model of recurrent positive selection [17]. If we found a significant difference between M7 and M8, then we tested for a difference between M8 and M8a, which approximates neutral evolution. We applied a Bonferroni correction for the M8-M8a p-value based on the number of genes that were run in the analysis. If the analysis for rejected the M7 null and the M8a null hypothesis, we then turned to a Bayes Empirical Bayes (BEB) analysis to identify amino acid positions that have statistical support for recurrent selection [74]. We considered a site to have strong evidence for recurrent selection if the BEB posterior probability was greater than 0.95. Finally, we checked that sites were not called as a result of poor alignment by checking for consistent alignment in the region surrounding each positively selected site.

### Whole genome scaling

Our analysis is designed to run on a gene by gene basis. To run the analysis for each gene in a reference genome, we broke the process into minimal computing elements and distributed the tasks over an array of computational resources using Amazon Web Services.

### Coincident recurrent selection analysis

To search for evidence of recurrent – recurrent selection, we limited our analysis to only genes that had a PAML result in all 6 of the clades we analyzed. We considered a gene to have strong evidence of recurrent selection only if there was at least one site with strong evidence for recurrent selection in that gene.

## Acknowledgments

This work was supported by the National Institutes of Health R01 GM115914 (NP), a Mario Capecchi endowed chair in Biology (NP), the Pew Biomedical Scholars Program (NP), and National Institutes of Health (Developmental Biology Training Grant 5T32 HD0741 (JCC).

## Supplemental Information

**Supplemental Figure 1.**
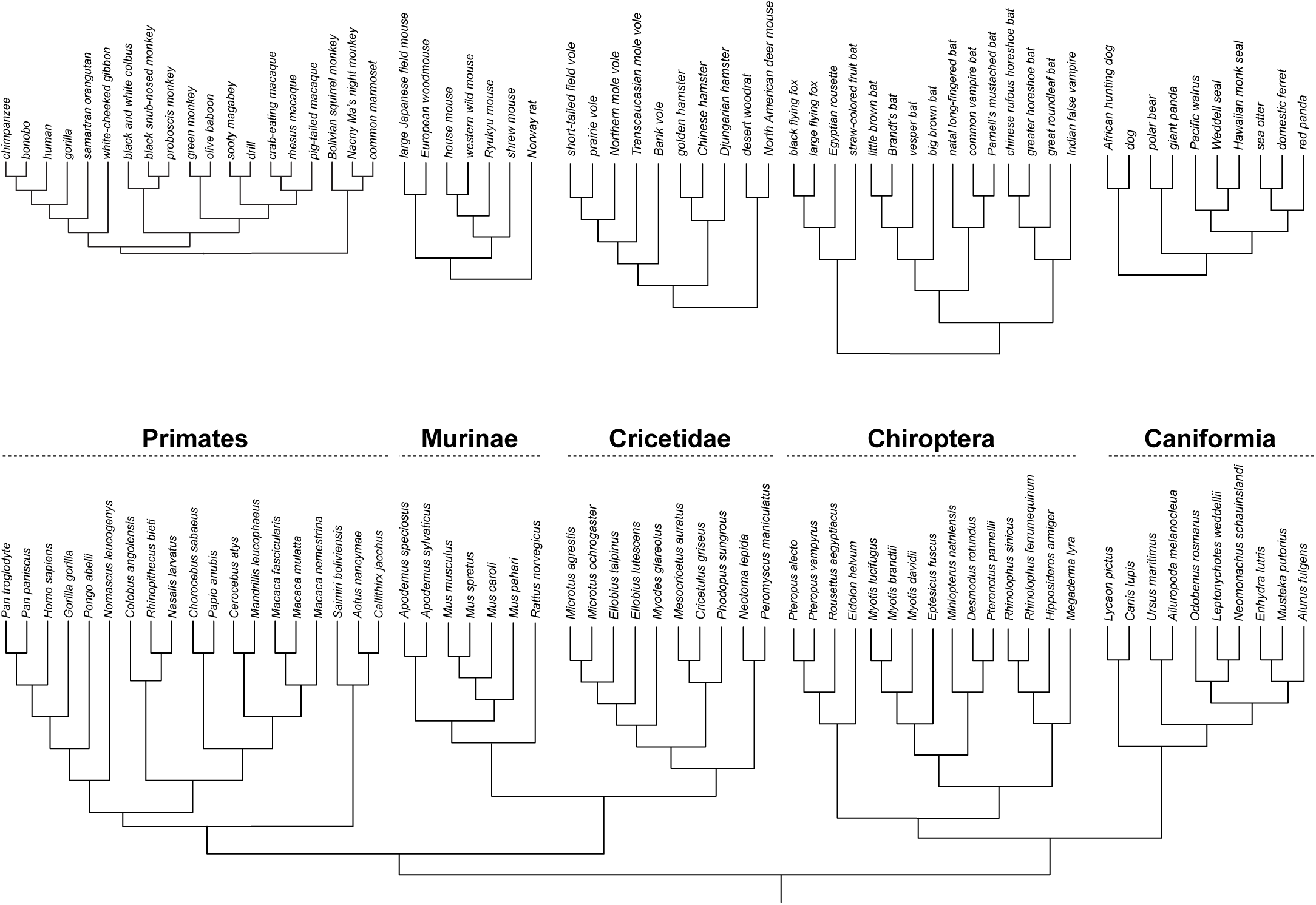
Phylogenies of mammalian clades used in this study. The taxonomic names are listed on the left for each species, with the common name listed on the right.

**Supplemental Figure 2.**
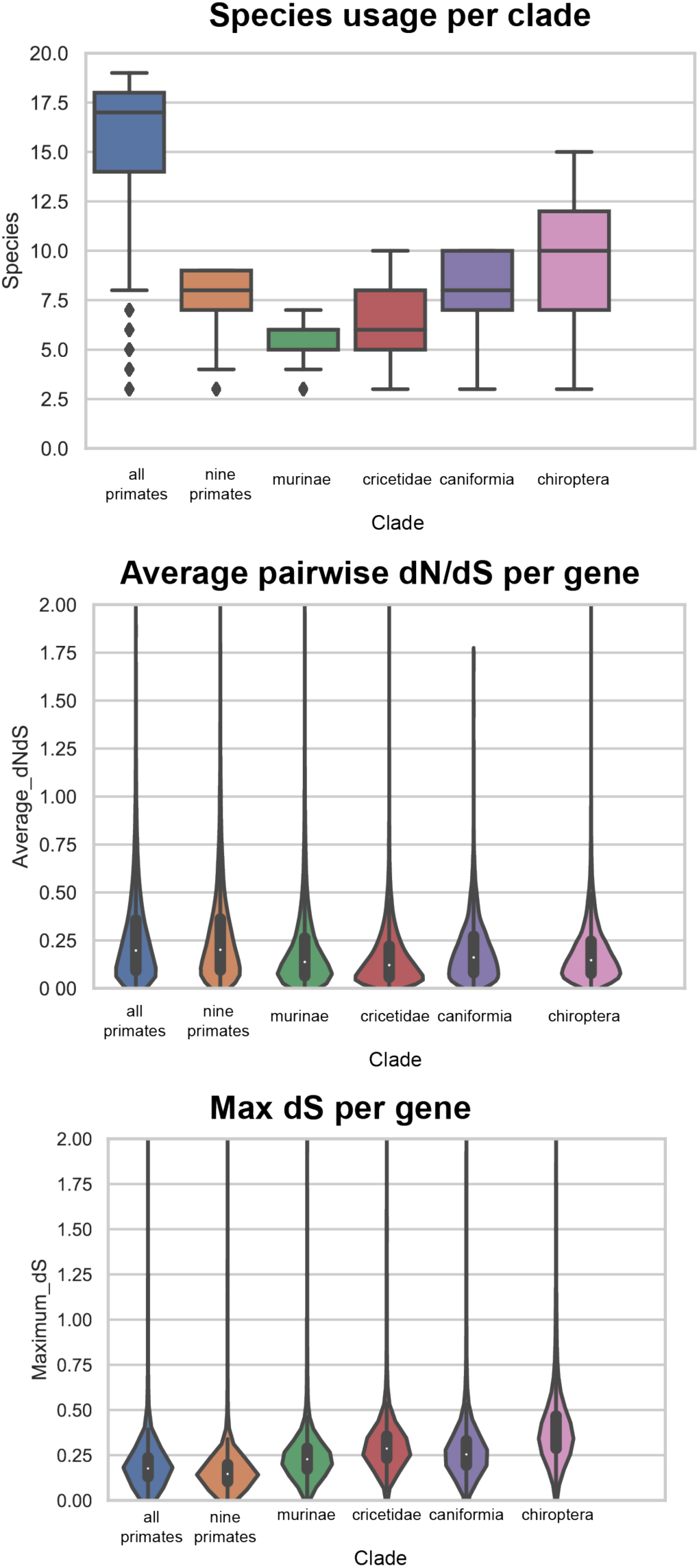
Summary statistics for PAML in mammalian clades.

**Supplemental Figure 3.**
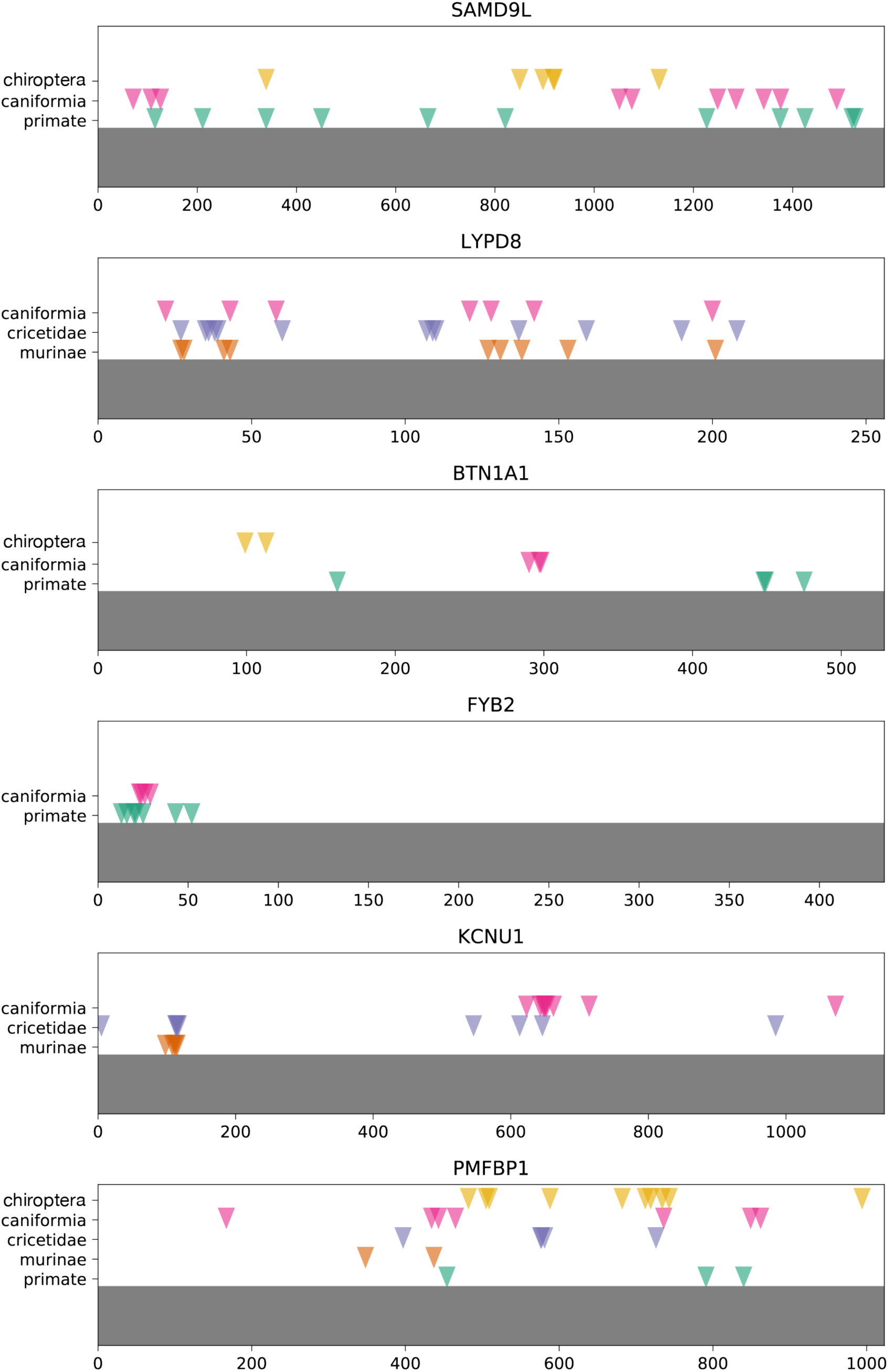
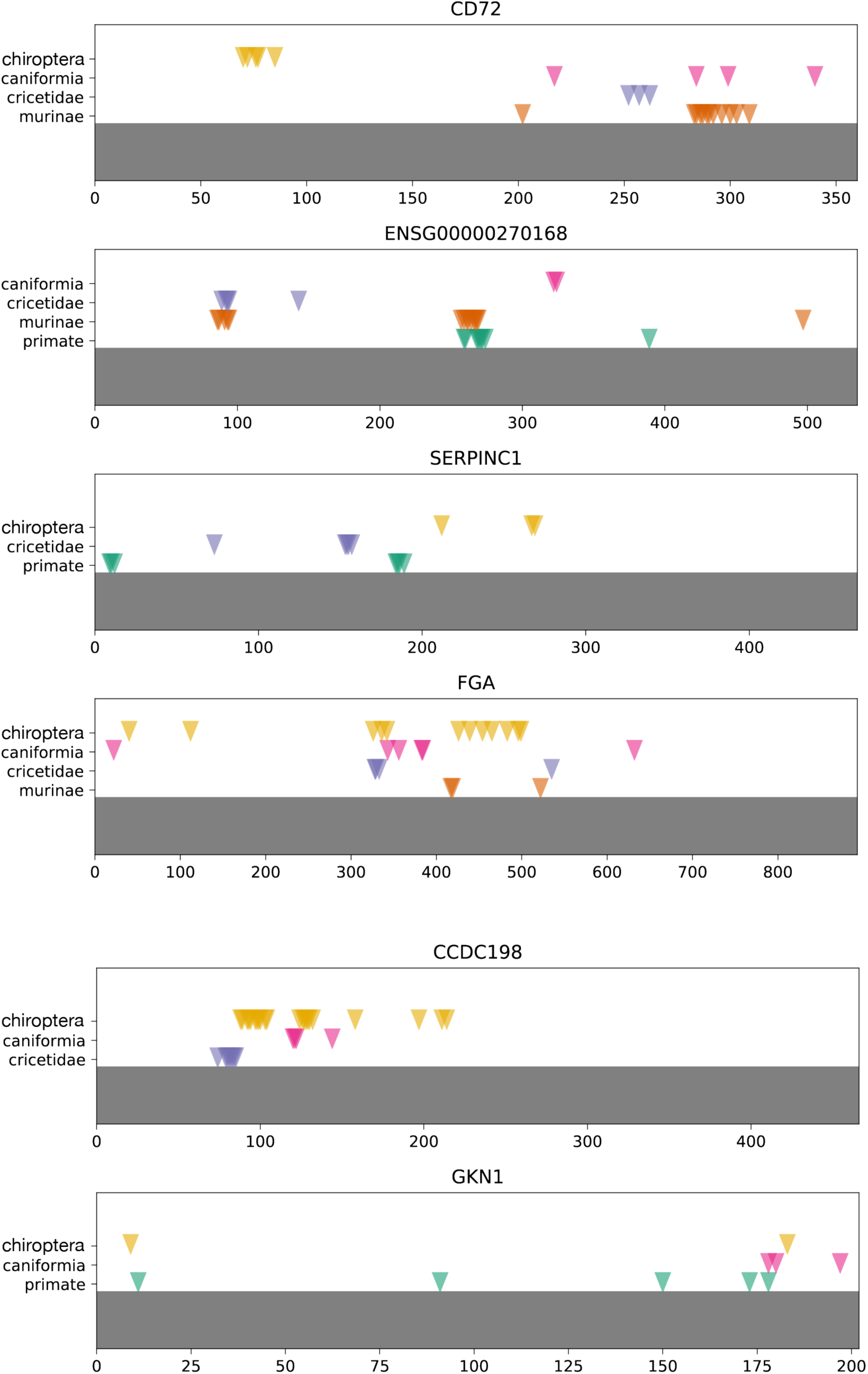
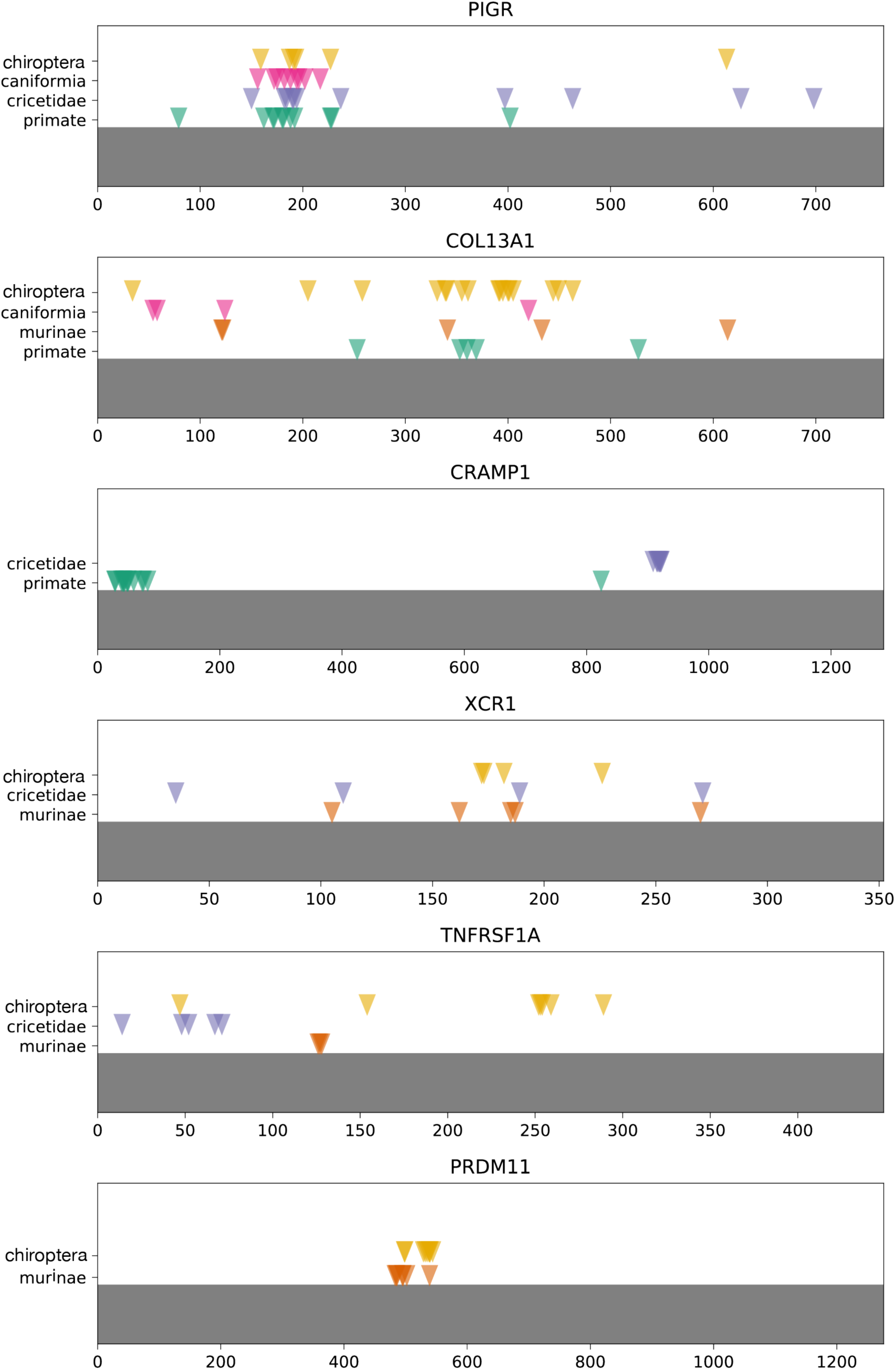
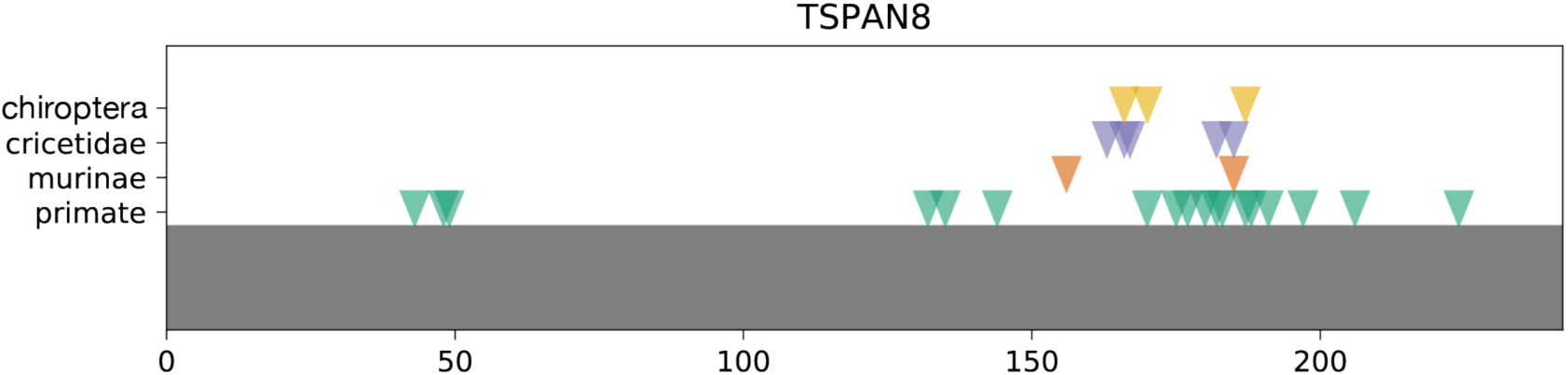
Sites under selection in genes with signatures of selection in three and four clades. The amino acid index of each gene is given on the x-axis. Sites under selection in each clade are denoted as triangles, with each clade on its own line.

## Supplemental Tables

**ST1 – Species and genomes used for this study.** Available online at bioRxiv.

**ST2 – Raw output of the computational pipeline.** Available online at bioRxiv.

